# Noncovalent antibody catenation on a target surface drastically increases the antigen-binding avidity

**DOI:** 10.1101/2022.07.12.499671

**Authors:** Jinyeop Song, Bo-Seong Jeong, Seong-Woo Kim, Seong-Bin Im, Wonki Cho, Myung-Ju Ahn, Byung-Ha Oh

**Author notes:** These authors contributed equally (Jinyeop Song, Bo-Seong Jeong). Department of Physics, Massachusetts Institute of Technology, Cambridge, MA, USA.

## Abstract

Immunoglobulin G (IgG) antibodies are widely used for diagnosis and therapy. Given the unique dimeric structure of IgG, we hypothesized that, by genetically fusing a homodimeric protein (catenator) to the C-terminus of IgG, reversible catenation of antibody molecules could be induced on a surface where target antigen molecules are abundant, and that it could be an effective way to greatly enhance the antigen-binding avidity. A thermodynamic simulation shows that quite low homodimerization affinity of a catenator, *e.g.* dissociation constant of 100 μM, can enhance nanomolar antigen-binding avidity to a picomolar level, and that the fold enhancement sharply depends on the density of the antigen. In a proof-of-concept experiment where antigen molecules are immobilized on a biosensor tip, C-terminal fusion of a weakly homodimerizing protein to two different antibodies enhanced the antigen-binding avidity by at least 210 to 5,120 folds from the intrinsic binding avidity. Thus, the homodimerization-induced antibody catenation would be a simple, powerful and general approach to improve many antibody applications, including the detection of scarce biomarkers and targeted anticancer therapies.

## INTRODUCTION

Immunoglobulin G (IgG) antibodies have become the principal therapeutic biologic. IgG antibodies are a homodimer of a heterodimer composed of two copies of each heavy chain (~50 kDa) and light chain (~25 kDa). They have two functional regions: the antigen-binding fragment (Fab) region at the N-terminal end and the fragment crystallizable (Fc) region at the C-terminal end. With an overall shape of the letter Y, the two identical regions of Fab form two arms that can bind two antigen molecules. This antibody-antigen engagement could prevent the antigen from binding to cognate partners or eliminate the antigen molecules from the cell surface by receptor-mediated endocytosis (Liu, 2018). The two copies of Fc form a homodimeric tail that enables a long half-life via binding to the neonatal Fc receptor (FcRn) and exerts effector functions via binding to the Fcγ receptors on effector immune cells or the complement factor C1q (Hogarth and Pietersz, 2012, Lee et al., 2017), which could lead to the death of cells to which antibody molecules are bound (Carter and Lazar, 2018, Goydel and Rader, 2021, Jiang et al., 2011).

IgG antibodies have desirable properties for use as a therapeutic drug, including high specificity for a target antigen, low immunogenicity and long serum half-life (Weiner et al., 2010). On the other hand, therapeutic monoclonal antibodies (mAbs) show side effects, albeit to a lesser degree in comparison with conventional chemotherapeutics, such as low or high blood pressure and kidney damage (Hansel et al., 2010). In the case of targeted cancer therapy, where mAbs target a specific antigen on cancer cells, the side effects likely arise due to the expression of the target antigen not only on cancer cells but also on normal cells, which therefore are targeted indiscriminately by mAbs administered in patients (Scott et al., 2012). Moreover, mAbs often suffer from shortcomings such as moderate therapeutic efficacy (resulting in the development of resistance) and their efficacy in a fraction of patients (as observed for mAbs against immune checkpoint inhibitors) (Aldeghaither et al., 2019, Hansel et al., 2010, Wang et al., 2021). Insufficient blockade of target antigens for various reasons, including insufficient antigen-binding affinity, could be responsible for the moderate therapeutic efficacy.

In general, diagnostic and therapeutic antibodies are required to exhibit low nanomolar or higher antigen-binding affinity (*K*_D_ < 10 nM) (Sliwkowski and Mellman, 2013). To reach this level of affinity, laborious experiments for affinity maturation are usually followed after an initial discovery of an antibody (Hoogenboom, 2005). Increasing the valency of binding interaction could be a method of choice. It was shown that irreversibly dimerized monovalent binders can bind targets significantly better than monomeric counterparts (Foreman, 2017). This enhancement of the binding affinity arises from the proximity effect, where the binding of one subunit of the dimer to a target restricts the search space for the other subunit. Reversibly dimerized binders could also exhibit significantly enhanced binding affinity depending on the affinity for binder-target interaction, affinity for homodimerization and the length of the connecting linker, as predicted by a reacted-site probability approach (Foreman, 2017). Such approaches to increase the valency of binding have been applied to IgG antibodies, and a considerable increase in the antigen-binding affinity was observed *in vitro* (White et al., 2014). However, as the size of an IgG-type antibody is large (~150 kDa), irreversible cross-linking or tight reversible dimer formation of the antibody would result in poor solubility and tissue penetration *in vivo*.

Owing to the overall dimeric structure, IgG antibodies genetically fused to a homodimeric protein at the C-terminus can be catenated in an arm-in-arm fashion as long as the homodimer can be formed, not within an antibody molecule, but between two antibody molecules. In theory, it would be possible to generate a soluble fusion protein that remains monomeric in solution, but becomes catenated by the proximity effect on a cell surface where target antigen molecules are abundant, provided that the fused protein has appropriately low homodimerization affinity. Importantly, this proximity effect-driven catenation, in turn, should result in enhanced bivalent antigen-binding affinity (=avidity). In this work, by agent-based modeling (ABM) and proof-of-concept experiments, we demonstrate that antibody catenation induced by the intermolecular homodimerization can enormously enhance the antigen-binding avidity of an antibody on a target surface.

## RESULTS

### The concept of antibody catenation on a target surface

This concept was based on (i) the unique dimeric structure of the IgG-type antibody and (ii) a proximity effect that potentially takes place on a target cell surface. In the structure of IgG, the Fc domain is composed of two copies of the constant regions of the heavy chain (C_H2_ and C_H3_) forming a homodimer, in which the two C-termini are ~23 Å apart and point away from each other (Figure 1A, *Left*). This structural feature indicated that a homodimer-forming protein genetically fused to the C-terminus can be prevented from forming a homodimer intramolecularly by controlling the length of the connecting linker or its homodimerization affinity. Instead, the fusion protein can form a homodimer intermolecularly, and then such a homodimerization could result in a catenation of the antibody molecules (Figure 1A, *Right*). We designate the fusion protein between an antibody and a homodimeric protein as antibody-catenator (^cat^Ab). A proximity effect for ^cat^Ab is expected on a target surface where multiple copies of target antigen are present, because the local concentration of ^cat^Ab on the surface will increase owing to the antibody-antigen binding interaction. Consequently, the homodimerization between the catenator molecules will increase to form catenated antibodies in an arm-in-arm fashion (Figure 1B). Importantly, the effective antigen-binding affinity of ^cat^Ab will increase in parallel with the catenation, and the fold enhancement would depend on the degree of the catenation. Thus, it appeared possible to enhance the antigen-binding avidity of the IgG-type antibodies by genetically fusing a weakly homodimer-forming protein.

**Figure 1.**
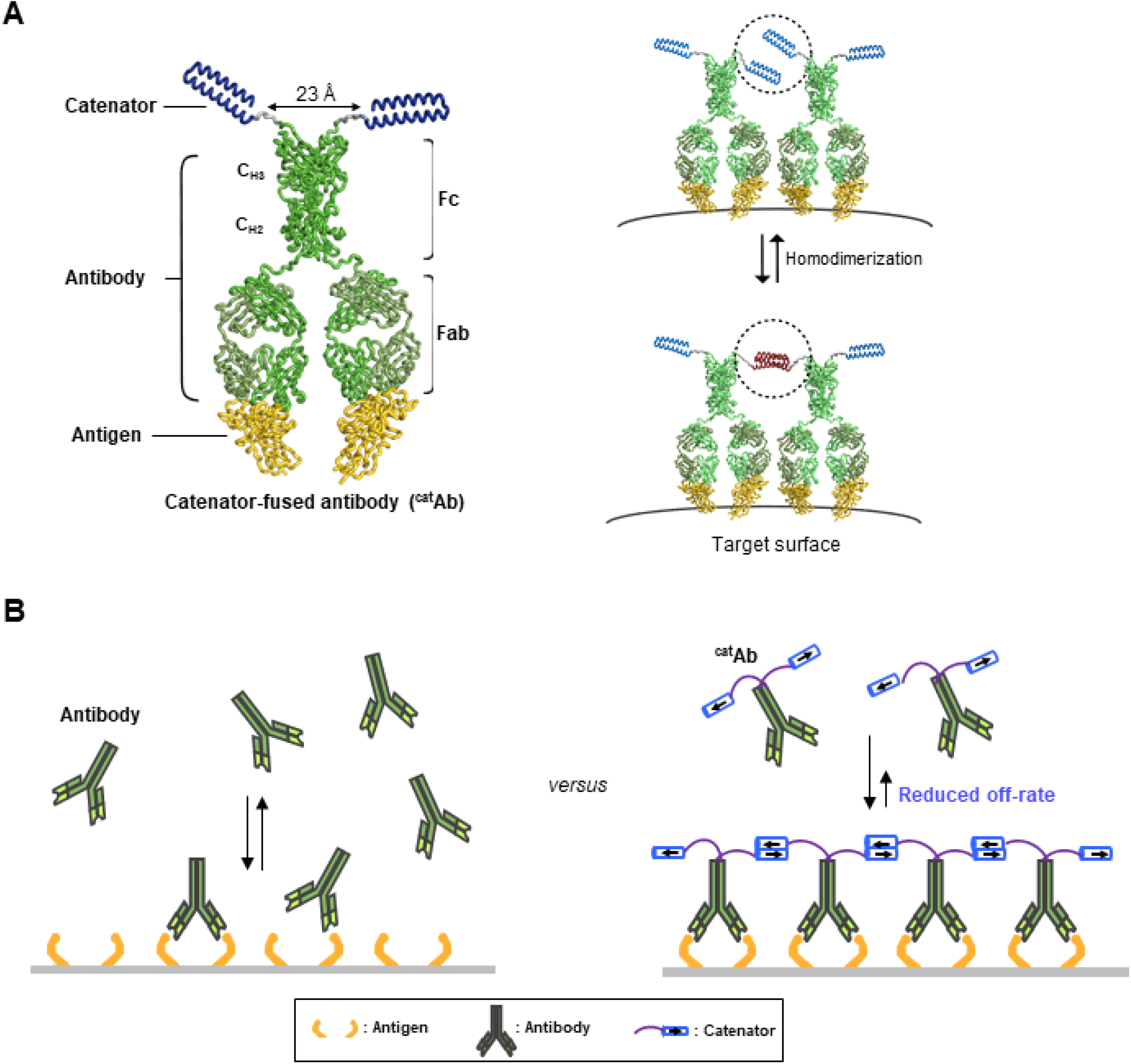
The concept of antibody catenation on a target surface by fusion of a catenator. (**A**) Molecular model for catenator-fused antibodies. A flexible linker (Gly-Gly-Ser) between Fc and the catenator and the hinge segment between Fc and Fab were modeled by using the ROSETTA software. The catenator is an α-helical hairpin that forms four-helix anti-parallel coiled coils (PDB entry: 1ROP). The structure of Fc was derived from the IgG1 antibody (PDB entry: 1IGY) and that of Fab from an antibody against the receptor-binding domain of the SARS-CoV-2 spike protein (PDB entry: 6XE1). (**B**) Decreased dissociation by antibody catenation. Pairs of ^cat^Ab-antigen complexes adjacent to each other can be catenated, and the ^cat^Ab molecules are increasingly harder to dissociate from each other with increased catenation. The effective antigen-binding avidity would increase owing to a decreased off rate of ^cat^Ab.

### Agent-based modeling to simulate the behavior of ^cat^Ab

ABM is a computational modeling approach that has been employed in a variety of research areas, including statistical physics (Perc et al., 2017, Fu and Wang, 2008) and biological sciences (An et al., 2009, Metzcar et al., 2019, McLane et al., 2011). ABM enables the understanding of macroscopic behaviors of a complex system by defining a minimal set of rules governing microscopic behaviors of agents which compose the system.

We constructed an ABM to simulate the behavior of the ^cat^Ab molecules on a target surface, where target antigen (Ag) molecules form antibody-binding sites. To circumvent complexity, we presumed that each binding site is a pair of two antigen molecules (2Ag), and ^cat^Ab make a bivalent interaction with the binding site in a 1:1 stoichiometry to form an occupied binding site (^cat^Ab-2Ag) (Figure 2A, *Left*). For catenation to occur between two adjacent ^cat^Ab-2Ag complexes, the distance between the centers of two adjacent complexes (*d*) should be closer than the reach length (*L*) defined as *l*+*c*/2, the sum of the linker length (*l*) and the half the catenator length (*c*) (Figure 2A, *Right*). Therefore, multiple parameters affect the catenation on the target surface. In our ABM model, we regarded every possible binding site on the target surface as an individual agent in the ABM formalism, and each binding site is assigned to a fixed position on a three-dimensional (3D) surface with a periodic boundary condition. Three rules in our ABM govern the behaviors of the ^cat^Ab molecules on the target surface. The first rule is about the *intrinsic antibody-antigen binding*. An unoccupied binding site binds to one free ^cat^Ab through bivalent interaction to form an occupied binding site. Bound ^cat^Ab may dissociate from the occupied binding site, leaving the binding site unoccupied. The equilibrium population of the occupied and unoccupied binding sites is determined by the antibody’s intrinsic avidity for the antigen with no effect of the catenator on the antigen-binding avidity assumed. Then, the relative likelihood of the occupied state compared to the unoccupied state for any binding site (the likelihood of intrinsic antigen binding) is defined as [^cat^Ab-2Ag]/[2Ag]) and thus can be expressed as [^cat^Ab]/*K*_D_, where [^cat^Ab] is the concentration of ^cat^Ab and *K*_D_ is the dissociation constant for the bivalent ^cat^Ab-2Ag interaction (Figure 2B, *Left*). The second rule is about *catenation*. A pair of ^cat^Ab-2Ag complexes on the target surface can be bridged by intermolecular homodimerization between catenators (Figure 2B, *Middle*). For a pair of ^cat^Ab-2Ag complexes separated by *d* (Figure 2A, *Right*), the relative likelihood of the catenation state as compared to the non-catenation state is the ratio of the forward reaction rate (catenation) to the reverse reaction rate (decatenation). The forward reaction rate (*R*_catenation_) and the reverse reaction rate (*R*_decatenation_) are given as,

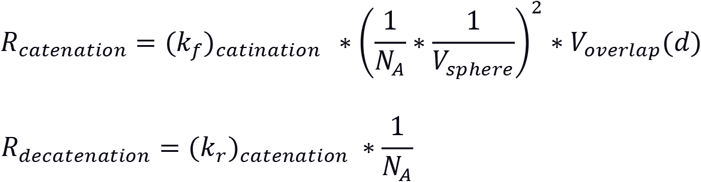
where *k*_f_ and *k*_r_ are the reaction rate constant of the forward and reverse reaction, respectively, *N_A_* is the Avogadro number, *V*_sphere_ is the local spherical volume within the reach of the catenator, and *V*_overlap_(*d*) is the volume where two catenators can come in contact to form a homodimer (Figure 2A, *Right*). In approximating the forward reaction rate, the catenator was assumed to sample *V*_sphere_ uniformly. The relative likelihood, defined as *R*_catenation_/*R*_decatenation_, is then expressed as

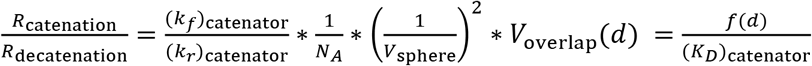

where

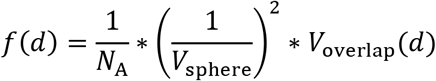

**Figure 2.**
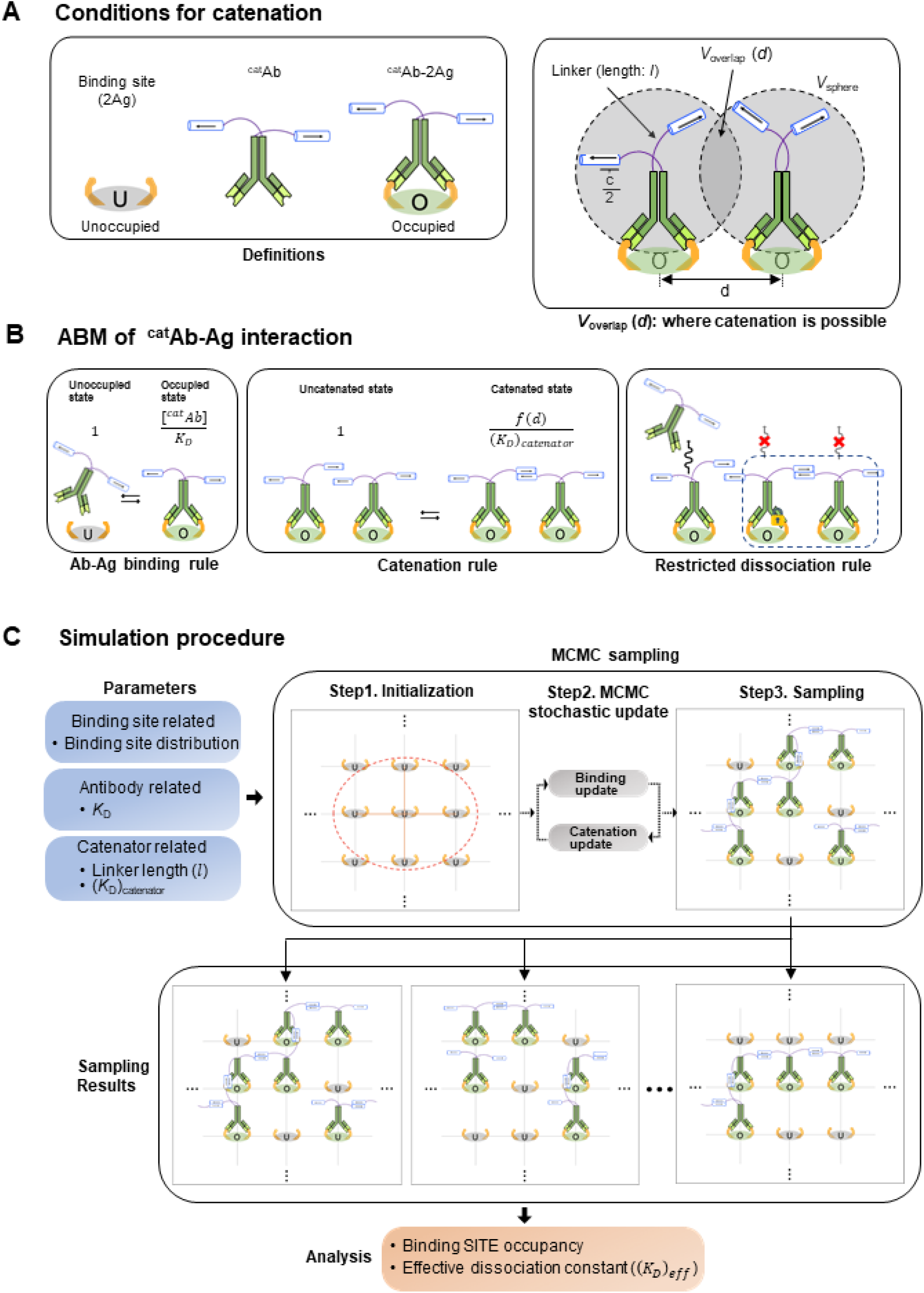
ABM for simulating the binding dynamics of a catenator-fused antibody. (**A**) (*Left*) Each binding site is composed of two antigen molecules (2Ag). (*Right*) The grey circles indicate the sphere sampled by the catenator, and *K*_overlap_ is the overlapping volume between the adjacent spheres. Catenation between two ^cat^Ab molecules is possible only in *V*_overlap_. (**B**) The three rules of the ABM model. (*Left*) ^cat^Ab-2Ag binding occurs with a relative likelihood, [^cat^Ab]/*K*_D_. (*Middle*) The catenation between adjacent ^cat^Ab-2Ag complexes occurs with an indicated relative likelihood, *f*(d)/(*K*_D_)_Catenator_, determined by (*K*_D_)_Catenator_ and the inter-complex distance *d*. (*Right*) It was assumed that ^cat^Ab molecules that are catenated cannot dissociate from the surface. (**C**) The simulation requires specification of the parameters for the binding site, antibody and catenator. Through the MCMC sampling, the state of binding sites on the target surface is iteratively updated with the ABM rules and eventually sampled. A sufficient number of sampling results are collected to quantify the binding occupancy and the effective dissociation constant.

The relative likelihood is thus a function of *d*, and it is inversely proportional to the dissociation constant of the catenator in the bulk medium, (*K*_D_)_catenator_. The function *f*(*d*) can be viewed as the effective local concentration of the catenator in *V*_overlap_(*d*). As expected, *f*(*d*) and thus the relative likelihood is sensitively affected by the reach length and limited by the ^cat^Ab-^cat^Ab distance (Figure 2—figure supplement 1). Finally, the third rule is about *restricted dissociation* which assumes that catenated antibodies are not allowed to dissociate from the binding site, because the catenated arms would hold the dissociated antibody near its binding site, forcing it to rebind immediately (Liese and Netz, 2018). Under this assumption, antibody molecules are allowed to dissociate from the binding site, only if its catenator is not engaged in the homodimerization with nearby ^cat^Ab-2Ag complexes (Figure 2B, *Right*).

### Simulations show significant enhancement of the antigen-binding avidity

According to the postulated rules, we simulated the effects of the antibody catenation on the binding interaction between ^cat^Ab and 2Ag on a three-dimensional surface by using the Markov Chain Monte-Carlo (MCMC) sampling method (Hooten and Wikle, 2010) (see Methods section). Our sampling procedure is composed of three steps (Figure 2C). The first step is an *initialization*, where a target surface with the antibody-binding sites is defined by specifying the coordinates for each site. A set of binding sites are positioned equidistant from each other or randomly positioned, and the inter-site distance or the number of binding sites were set as variables. The next step is an *MCMC stochastic update* step. In each iteration, a binding site is randomly selected from the target surface, and the probability of changing the status of the selected binding site (occupied or not) is calculated by the Metropolis-Hasting algorithm (Hastings, 1970, Grazzini et al., 2017). Then, the ‘on’ or ‘off’ state of this site is updated with the calculated probability. Accordingly, the catenation state is probabilistically updated for each update step. In the following sampling step, the total number of the occupied binding sites is counted, which is then collected through multiple simulation runs for the statistical analysis of the binding site occupancy and the effective antigen-binding avidity. The binding site occupancy is the mean value of the number of occupied binding sites collected for more than 1024 MCMC samplings. For each simulation, we calculated the mean binding occupancy and the effective dissociation constant, (*K*_D_)_eff_, which takes into account the effect of the antibody catenation, and is expressed as:

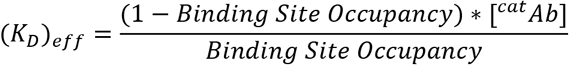

Since the catenator homodimerization should be affected by how the binding sites are distributed on a 3D surface, simulations were conducted for different arrays of binding sites. In the simulations, (*K*_D_)_catenator_ was the main variable, while other parameters were set constant. First, we simulated the binding sites forming a square lattice to find that the Ab-catenator exhibited enhanced binding site occupancy in a sigmodal manner, and that it could be enhanced to near full saturation by a catenator that forms a homodimer with quite low binding affinity. For instance, a ^cat^Ab with (*K*_D_)_catenator_ of ~1 μM exhibited ~70-fold enhancement of the effective antigen-binding avidity (=reduction of (*K*_D_)_eff_) in comparison with the same antibody without a fused catenator (Figure 3). As a means of comparison across different simulation setups, we employed ‘(*K*_D_)_catenator,50_’ which is defined as the (*K*_D_)_catenator_ that enables half-maximal enhancement of the binding site occupancy (Figure 3).

**Figure 3.**
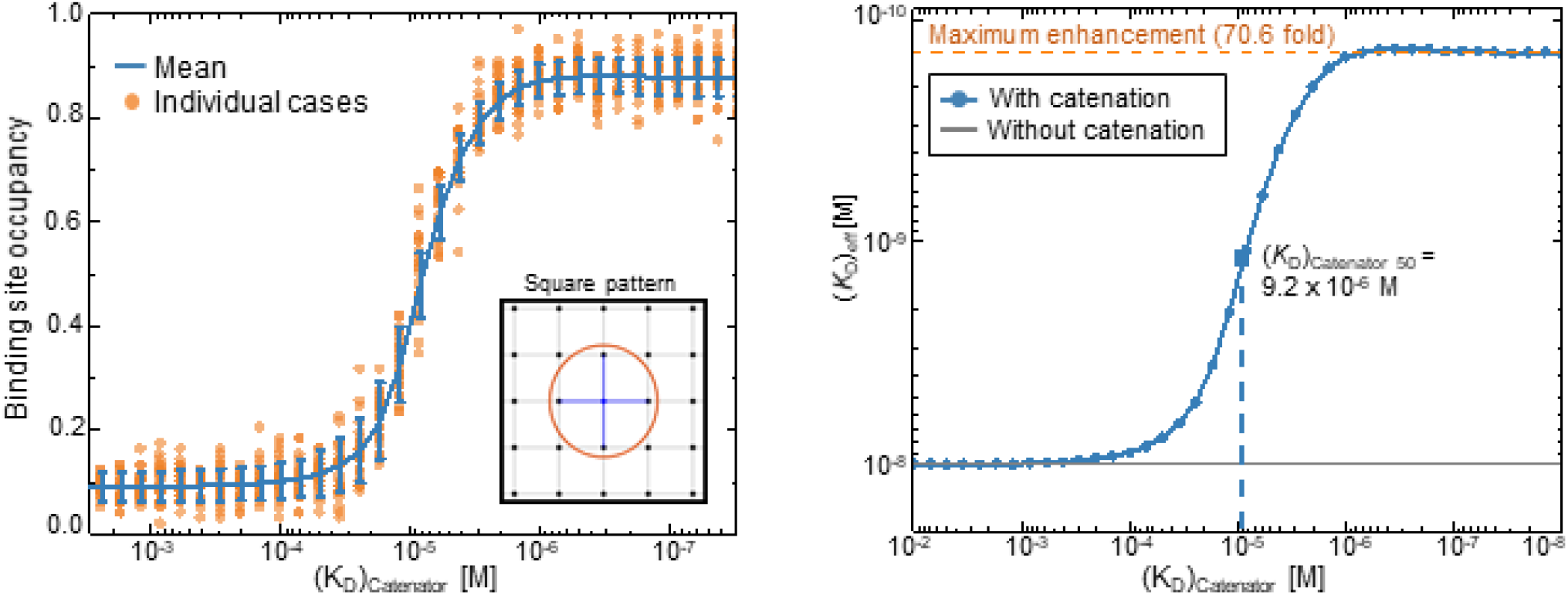
Simulations of the binding site occupancy and (*K*_D_)_eff_ in response to (*K*_D_)_catenator_. (*Left*) Binding site occupancy. The simulations were carried out for a square array of the binding sites. The values for a set of variables were *K*_D_ = 10^-8^ M, [^cat^Ab]= 10^-9^ M, reach length= 7 nm, spacing between the binding sites= 12 nm and the number of total binding sites= 98. The mean value and standard deviations of 1024 MCMC simulations for each (*K*_D_)_catenator_ value are shown in blue, and the data are shown as a scatter plot of representative runs (orange). (*Right*) The effective dissociation constant. The data shown on the left were converted into the (*K*_D_)_eff_ values. The dashed line represents the *K*_D_ value for the same antibody without a catenator. The maximum fold enhancement of the effective binding avidity, which is equivalent to the reduction of (*K*_D_)_eff_, is 70.6.

### Comparison of the simulations for different arrays of the binding sites

Next, we carried out simulations for other regular arrays of the binding sites and for randomly distributed binding sites. Depending on the pattern of regularly distributed binding sites, the number of possible catenations for a given binding site (designated as connectivity number) varies: 3, 4 and 6 for a hexagonal, square or triangular array of the binding sites, respectively (Figure 4A). These three arrays showed varying but similar enhancement of the binding site occupancy and the effective antigen-binding avidity by the catenator (Figure 4A). As expected, the higher the connectivity number was, the lower (*K*_D_)_catenator,50_ an array exhibited; the (*K*_D_)_catenator,50_ was 8.0, 9.2 and 12.2 μM for the hexagonal, square and triangular array of the binding sites, respectively. The simulations showed that, as the connectivity number increased, the effective antigen-binding avidity increased with the maximum 41-, 73- and 93-fold enhancement for the triangular, square and hexagonal array, respectively (Figure 4A). Thus, regardless of the distribution patterns, the effective antigen-binding avidity could be increased by at least 41 folds under the simulation conditions where the target surface contains only 98 binding sites.

**Figure 4.**
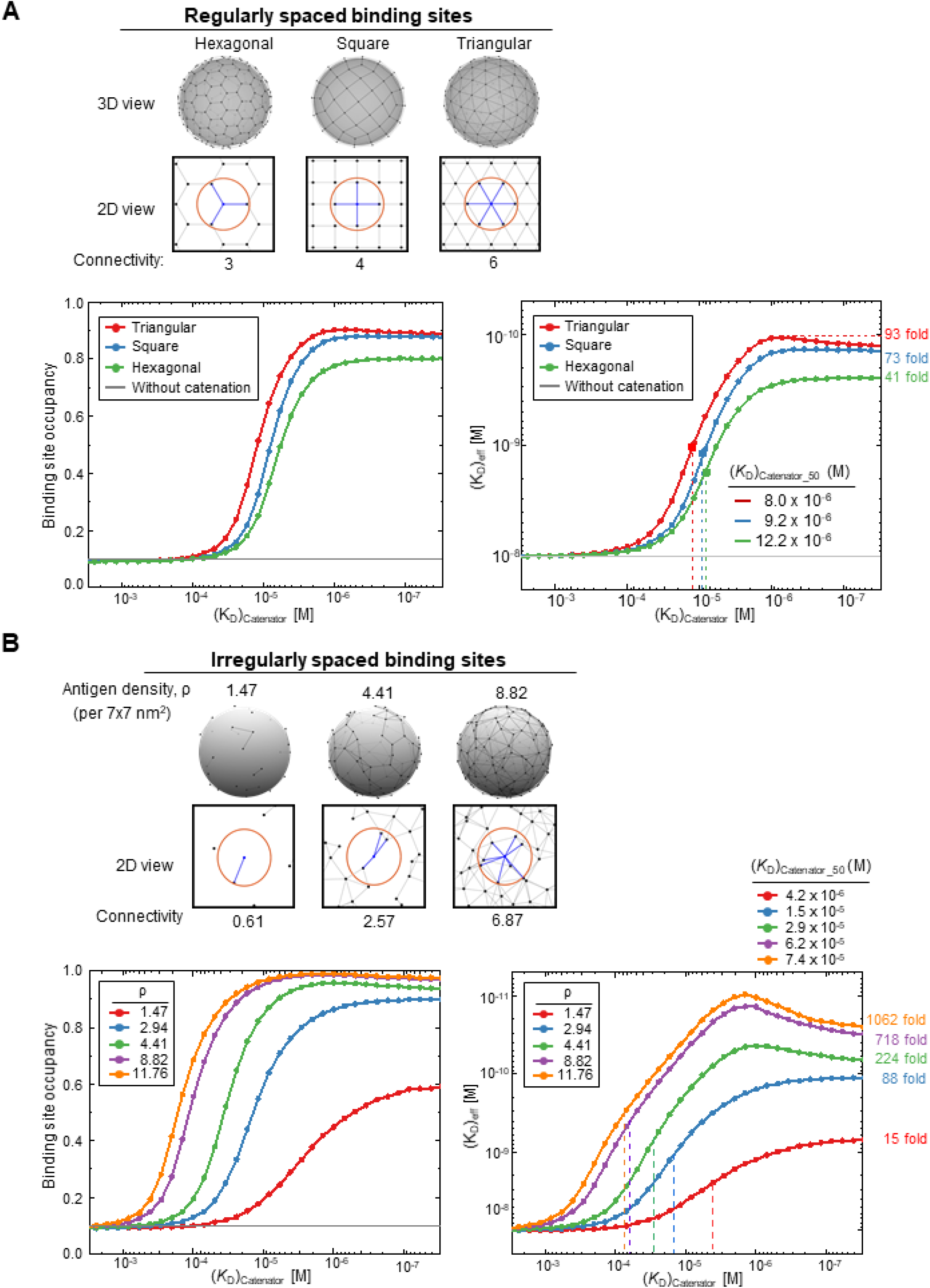
Simulations for different arrays of the binding sites. (**A**) Comparison for regularly distributed binding sites. Three different regular arrays of the binding sites are shown at the top. The black dots represent the binding sites and grey lines the connectable pairs by the catenators. The red circles and the blue lines represent the maximum range of catenation and the connectivity number, respectively, for a given binding site. Binding site occupancy and (*K*_D_)_eff_ in response to (*K*_D_)_catenator_ are shown at the bottom. 1024 trials were sampled for each (*K*_D_)_catenator_ value and the results are plotted. The variables were *K*_D_= 10^-8^ M, [^cat^Ab]= 10^-9^ M, reach length= 7 nm, spacing between the binding sites= 12 nm, and the number of total binding sites were 98 for the square array and 102 for hexagonal and triangular array, respectively. The numbers on the right are the maximum fold enhancement of the effective binding avidity for each array. (**B**) Comparison for randomly distributed binding sites. Three random arrays of the binding sites with different binding site density (ρ) are shown at the top. The surface area for the simulation was 5,760 nm^2^. The simulation conditions were the same as in (A). Binding site occupancy and (*K*_D_)_eff_ in response to (*K*_D_)_catenator_ are plotted as in (A).

For the case of randomly distributed binding sites on a 3D surface, which is relevant to target antigen distribution on cell surfaces, we introduced the binding site density (ρ), the number of binding sites per unit area which is set to the square of the reach length (7 nm) (Figure 4B). In the simulations, the total surface area was 5,760 nm^2^, and the number of binding sites was 15, 30, 45, 90 or 120, which correspond to the ρ of 1.47, 2.94, 4.41, 8.82 or 11.76. Denser binding sites would increase the connectivity number for a given binding site. As expected, simulations showed that higher binding site density resulted in a higher level of binding site saturation and a much more significant increase in the effective antigen-binding avidity; the maximum fold enhancement ranged from 15 (ρ= 1.47) to 1,062 (ρ= 11.76). Likewise, significantly different (*K*_D_)_catenator,50_ values were observed: *e.g*., 4.2×10^-6^ M at the ρ of 11.76 *versus* 74×10^-6^ M at the ρ of 1.47 (Figure 4B). The maximal saturation and the onset (*K*_D_)_catenator_, which begins to exert the catenation effect, were also considerably different. Thus, the catenation effects are sensitively affected by the binding site density, in contrast with the all-or-none catenation effect observed for the regular arrays of the binding sites (Figure 4B).

In particular, the catenation-induced enhancement of the antigen-binding avidity was remarkably and sensitively affected by the (*K*_D_)_catenator_ values at high binding site density (ρ> 4.41) (Figure 4B). Much greater enhancement was observed as we further increased the density of randomly distributed binding sites: ~29,000 maximum fold enhancement at the ρ of 58.8 (Figure 4—figure supplement 1), which roughly corresponds to two hundredths of the density of the HER2 receptor on HER2-overexpressing breast cancer cells (Peckys et al., 2019). Together, our simulations show that randomly distributed binding sites at high density enormously enhance the effective antigen-binding avidity of ^cat^Ab.

Additionally, we performed simulations for different values of [^cat^Ab]/*K*_D_ to estimate the effect of *K*_D_ with respect to [^cat^Ab]. Varying [^cat^Ab]/*K*_D_ from 0.01 to 1.0 resulted in 85- to 900-fold enhancement of the antigen-binding avidity, suggesting that the catenation effect works for a broad range of *K*_D_ values (Figure 4—figure supplement 2).

### Proof-of-concept experiments

For experimental validation, we chose stromal cell-derived factor 1α (SDF-1α) as a catenator. SDF-1α is a small (*Mr*= 8 kDa) and weakly homodimerizing protein (*K*_D_= 150 μM) (Veldkamp et al., 2005), indicating that this protein fused to an antibody by a ~40 Å-long linker would not form an intramolecular homodimer within a fusion protein. By using a 10-residue connecting linker (GGGGSGGGSGG), SDF-1α was fused to two different antibodies: Trastuzumab(N30A/H91A), a variant of the clinically used anti-HER2 antibody Trastuzumab and glCV30, an antibody against the receptor-binding domain (RBD) of the severe acute respiratory syndrome coronavirus 2 (SARS-CoV-2) spike protein. The Fab fragment of Trastuzumab(N30A/H91A) binds to the ectodomain of HER2 with a *K*_D_ of 353 nM (Slaga et al., 2018), and glCV30 binds to the RBD with a similar binding affinity (*K*_D_= 407 nM) (Hurlburt et al., 2020). We produced the SDF-1α-fused antibodies, Trastuzumab(N30A/H91A)-SDF-1α and glCV-SDF-1α, and also the unmodified antibodies to compare their binding avidities by bio-layer interferometry (BLI) where respective target antigen was immobilized on a sensor tip. Unlike the similar antigen-binding affinities of the Fab fragments, the binding avidities of the full-form antibodies were quite different in our quantification, Trastuzumab(N30A/H91A) and glCV30 exhibiting the *K_D_* of 2.1 nM and 51.2 nM, respectively (Figure 5). The SDF-1α-fused antibodies exhibited association kinetics similar to those of the mother antibodies; association rate constants (*k*_aS_) of the Trastuzumab(N30A/H91A) and Trastuzumab(N30A/H91A)-SDF-1α were 1.5×10^5^ Ms^-1^ and 3.1×10^5^ Ms^-1^, respectively. Similarly, those of glCV30 and glCV30-SDF-1α were 25.0×10^4^ Ms^-1^ and 5.0×10^4^ Ms^-1^, respectively. However, the two pairs of the antibodies exhibited significantly different dissociation kinetics; dissociation rate constants (*k*_dS_) of the Trastuzumab(N30A/H91A) pair were 3.1×10^-4^ Ms^-1^ *versus* < less than 1.0×10^-7^ Ms^-1^ and those for the glCV30 pair were 1.3×10^-4^ Ms^-1^ *versus* less than 1.0×10^-7^ Ms^-1^ (Figure 5). These observed kinetics are consistent with the expectation that fused SDF-1α would not affect the association of the antibodies, but would slow down the dissociation of the SDF-1α-fused antibodies into the bulk solution, as it catenates the antibody molecules on the sensor tip. As a result, Trastuzumab(N30A/H91A)-SDF-1α exhibited the *K*_D_ of < 10 pM, at least 210-fold higher binding avidity compared with Trastuzumab(N30A/H91A), and likewise, The SDF-1α fusion to glCV30 increased the binding avidity by at least 5,120 folds, demonstrating that two-digit nanomolar binding avidity of an antibody can be increased to picomolar binding avidity by fusing a weakly homodimerizing protein.

**Figure 5.**
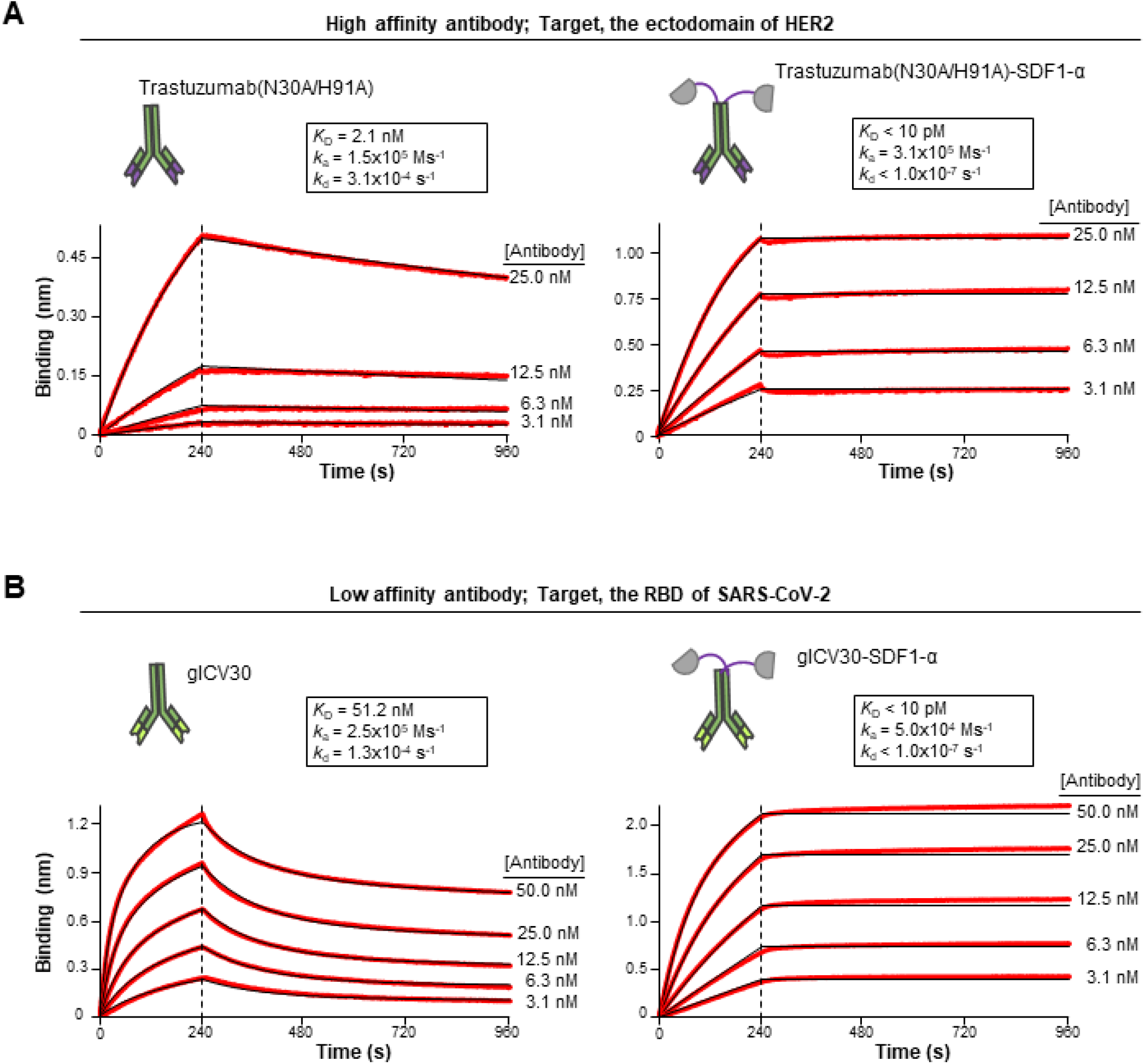
BLI runs demonstrating the effect of catenation on the binding avidity. The binding kinetics were measured with the indicated targets immobilized on a sensor tip. The concentration of the antibodies was varied as shown. The experimental signals and fitted curves are shown in red and black, respectively. For curve fitting, 1:1 binding was assumed. The kinetic parameters are shown in the insets. *k*_a_, association rate constant; *k*_d_, dissociation rate constant. (A) High-affinity antibody. Trastuzumab(N30A/H91A) exhibited the *K*_D_ of 2.1 nM for the immobilized ectodomain of HER2. (B) Low-affinity antibody. glCV30 exhibits the *K*_D_ of 51.2 nM for the immobilized RBD of SARS-CoV-2. For both Trastuzumab(N30A/H91A)-SDF-1α and glCV30-SDF-1α, the *K*_D_ values could not be accurately determined due to the instrumental insensitivity (*K*_D_ < 10 pM). The experiments were performed in triplicates, and representative sensorgrams are shown.

## DISCUSSION

In the current phage display for antibody screening, many candiates that do not satisfy a required affinity for a target antigen are rejected, although they might have high specificity of binding. A simple and general way of increasing the antigen-binding affinity of antibodies would be highly valuable for various applications of antibodies. Taking advantage of the particular homodimeric structure of IgG antibodies, we put forth a concept to enhance the bivalent antigen-binding interaction by fusing a weakly homodimerizing protein to the C-terminus of Fc. The validity of the concept was tested by simulations based on an ABM and supported by experimental demonstrations.

Our ABM with the three postulated rules was the basis for predicting the enhancement of effective antigen-binding avidity. The model has caveats. First, the assumption of uniform density for the fused catenators within a sphere oversimplifies the dynamics of the catenators, which would highly depend on physical contexts, such as molecular orientations and potential intramolecular interaction with the antibody (Zhou, 2001). Second, the binding sites representing antigens are fixed on a surface in our model, but in real situations, antigens move their positions, *e.g*., receptor molecules on cellular membranes (Saxton and Jacobson, 1997). Advanced molecular dynamics simulations incorporated into the ABM would take account of these microscopic details to result in a more accurate prediction of the behaviors of the ^cat^Ab molecules and the binding sites. Despite these caveats, the simulations provided valuable insights into the proper ranges of the antigen-binding avidity of an antibody and catenator-catenator binding affinity. According to the simulation, we adopted SDF-1α, a weakly homodimerizing protein (*K*_D_ of 150 nM), as a catenator. When fused to antibodies, it resulted in remarkable enhancement of the effective antigen-binding avidity of the antibodies, which was due to drastically reduced rate of dissociation of the fused antibody molecules from the immobilized antigens.

The “antibody catenation on a target surface” method presented herein might find practical applications. First, it can be applied to therapeutic antibodies against viruses, which have multiple copies of target antigens on their surface. Second, it can be used for sandwich-type point-of-care biosensors in which a second antibody is catenated to increase the sensitivity of detection. Third, this method can be used to sense biomarkers that exist in a very low number on a target cell (*e.g*., copy number < 10), which requires an extremely high-binding avidity of a probe antibody. For this application, employing an antibody with high antigen-binding affinity (*e.g*., *K*_D_ < 1 nM) and a catenator with high homodimerization affinity (*e.g*., (*K*_D_)_catenator_ < 1 μM), would be necessary to overcome low proximity effect due to the scarcely present antigen molecules. Fourth, it might also be applied to antibody-based targeted cancer therapy where side effects arising from antibody binding to normal cells are a general problem. Since cancer-associated antigen molecules are lower in number on normal cells than they are on cancer cells, catenated anticancer antibodies would be concentrated on the surface of cancer cells, because the effective binding avidity of a ^cat^Ab depends on the number of antigen molecules on a target surface. In particular, this approach would greatly reduce the intrinsic toxicity of antibody-drug conjugates that are widely used currently. Of note, a catenator fused to the C-terminus of Fc would not affect the effector functions of Fc through the Fcγ receptor-binding site and FcRn binding site on it. We observed that ^cat^Ab molecules could be internalized into HER2-positive cells presumably via receptor-mediated endocytosis (data not shown).

In conclusion, the presented strategy of the antibody catenation on a target surface is simple and powerful, and thus it could be widely applicable, although the homodimerization affinity of a catenator and the length of the linker need to be optimized case by case. Improvement of the simulation method will better guide the decision on the variables in constructing catenator-fused antibodies.

## Materials and Methods

### MCMC simulation

Simulation runs were carried out in the three steps stated below with specification of the target surface, *K*_D_ (for antibody-antigen interaction), (*K*_D_)_catenator_ (for catenator-catenator interaction) and *f*(*d*) (effective local concentration of the catenator). In all simulations, the number of ^cat^Ab was far more than that of the binding sites, and therefore, the concentration of free ^cat^Ab was assumed to be the same as that of total ^cat^Ab (free ^cat^Ab + antigen-bound ^cat^Ab). Simulation parameters their set values are listed in Table 1.

**Table 1.** Simulation specifications. The definition and values of the parameters used in the presented simulations are tabulated:

| Parameters | Description | Values |
| --- | --- | --- |
| **Specification of <sup>cat</sup>Ab** |  |  |
| $K_D$ | Dissociation constant of antibody | 10 nM |
| $(K_D)_{\text{catenator}}$ | Dissociation constant of catenator | 10 nM-10 mM |
| $[\text{catAb}]$ | Antibody concentration | 1 nM |
| $l$ | Length of the flexible linker | 6 nm |
| $c$ | Length of the catenator | 2 nm |
| $L$ | Reach length ($l+c/2$) | 7 nm |
| **Specification of the target surface** |  |  |
| $N_{\text{total\_binding\_sites}}$<br>(in Fig. 3,4) | Number of antibody-binding sites | 98 - 102 |
| Connectivity number<br>(in Fig. 3,4) | Number of possible catenation | 3 (Hexagonal)<br>4 (Square)<br>6 (Triangular) |
| $d$ (in Fig. 3,4) | Distance between adjacent binding sites | 12 nm |
| $L_{\text{surface}}$ (in Fig. 4) | Surface area of the target surface | 40 nm<sup>2</sup> |
| Binding site density<br>(in Fig. 4) | Surface density of the binding sites | 1.47-11.76<br>(per 7x7 nm<sup>2</sup>) |
| **Specification of Simulation** |  |  |
| Updates/MCMC step | Number of updates in one MCMC step | 30,000-100,000 |
| Sampling size | Number of sampling for a parameter set | 1,024 |

#### Step 1. Initialization step

1. A specified 3D target surface is implemented by assigning binding sites to specific locations on the surface.
2. Each binding site is set to be unoccupied.

#### Step 2. *MCMC stochastic update* step

The following sub-steps (1-3) are iterated sufficient times to ensure thermodynamic equilibration.

1. A random binding site *BS1* is chosen from the target surface.
2. The binding status of *BS1* is updated. If *BS1* is unoccupied, its status is changed to the occupied status with the acceptance probability of 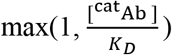. If *BS1* is occupied, its status is changed to the unoccupied status with the acceptance probability of 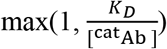).
3. An occupied binding site *BS2* right next to *BS1* is picked on the target surface, and the catenation status of the pair (*BS1, BS2*) is updated. If (*BS1, BS2*) is uncatenated, and if both *BS1* and *BS2* have an unengaged catenator, its status is changed to the catenation status with the acceptance probability of 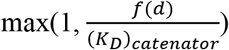. If (*BS*_1_, *BS*_2_) is catenated, its status is changed to the uncatenated status with the acceptance probability of 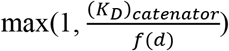

#### Step 3. Sampling step

1. The update step is stopped, and the final status of the target surface is recorded.
2. The total number of occupied and unoccupied binding sites are counted.

The codes for the model system and simulations are available in MATLAB and available on Github (https://github.com/JinyeopSong/Antibody_ThermoCalc_JY). The detailed description is provided in Readme.

### Preparation of antibodies and catenator-fused antibodies

Each DNA fragment encoding heavy chain variable regions (V_H_) and light chain variable regions (V_L_) of glCV30 were synthesized (IDT) and cloned into the pCEP4 vector (Invitrogen). DNA fragments of C_H1_-C_H2_-C_H3_ of the gamma heavy chain and C_L_ of the kappa-type light chain were inserted into the V_H_ and V_L_, and the resulting vectors were named glCV30 Hc and glCV30 Lc, respectively. DNA fragment encoding SDF-1α was synthesized (IDT) and cloned into the glCV30 Hc next to C_H3_ of glCV30 with (Gly-Gly-Gly-Gly-Ser)_2_ linker sequence (glCV30-SDF-1α Hc). For antibody production, the three vectors were amplified using the NucleoBond Xtra Midi kit (Macherey-Nagel), and a combination of the glCV30 Hc and glCV30 Lc vectors or a combination of the glCV30-SDF-1α Hc and glCV30 Lc vectors were introduced into the CHO-S cells (Gibco). The transfected cells were grown in the ExpiCHO expression medium (Gibco) for ten days post-transfection. Supernatants were collected by centrifugation at 4 °C, filtered through 0.45 μm filters (Millipore), diluted by the addition of a binding buffer (150 mM NaCl, 20 mM Na_2_HPO_4_, pH 7.0) to a 1:1 ratio, loaded onto an open column containing Protein A resin (Sino Biological), and eluted with an elution buffer (0.1 M glycine, pH 3.0). The eluent was immediately neutralized by a neutralizing buffer (1M Tris-HCl, pH 8.5), and the antibodies were further purified using a HiLoad 26/60 Superdex 200 gel-filtration column (Cytiva) equilibrated with a buffer solution containing 20 mM Tris-HCl (pH 7.5) and 150 mM NaCl. For preparing Trastuzumab(N30A/H91A) and Trastuzumab(N30A/H91A)-SDF-1α, each DNA fragment encoding V_H_ and V_L_(N30A/H91A) of Trastuzumab was synthesized (IDT). The cloning, protein production and purification procedures were virtually identical to those used for glCV30 and glCV30-SDF-1α.

### Bio-layer interferometry

BLI experiments were performed to measure dissociation constants using an Octet R8 (Sartorius). Biotinylated SARS-CoV-2 RBD (Acrobio system) or biotinylated Her2/ERBB2 (Sino Biological) was loaded to a streptavidin biosensor tip (Sartorius) for 120 s. A baseline was determined by incubating the sensor with Kinetics Buffer (Sartorius) for 60 s. Antibody samples at different concentrations went through the association phase for 240 s and the dissociation phase for 720 s. All reactions were carried out in the Kinetics Buffer (Satorius). The binding kinetics were analyzed using the Octet DataAnalysis 10.0 software (Sartorius) to deduce the kinetic parameters. Experiments were performed in triplicate for glCV30 and glCV30-SDF-1α and duplicate for Trastuzumab(N30A/H91A) and Trastuzumab(N30A/H91A)-SDF-1α.

## Figure preparation

The computational models of an antibody and an antibody-catenator in Figure 1A were generated by using the ROSETTA software (Leman et al., 2020), and are presented by PyMOL (DeLano, 2004).

## ACKNOWLEDGMENTS

This work was supported by the Samsung Research Funding & Incubation Center of Samsung Electronics under Project Number SRFC-MA2002-06.

## AUTHOR CONTRIBUTIONS

B.-H.O. directed the work, and B.-S.J., J.S., W.C. and M.-J.A. further conceptualized the research. J.S. performed simulations. B.-S.J., S.-W.K. and S.-B.I. performed cloning and purification of the antibodies and the SARS-CoV-2 RBD. B.-H.O., J.S., B.-S.J. and W.C. wrote the original draft. All authors reviewed and accepted the final manuscript.

## Figure supplements

**Figure 2—figure supplement 1.**
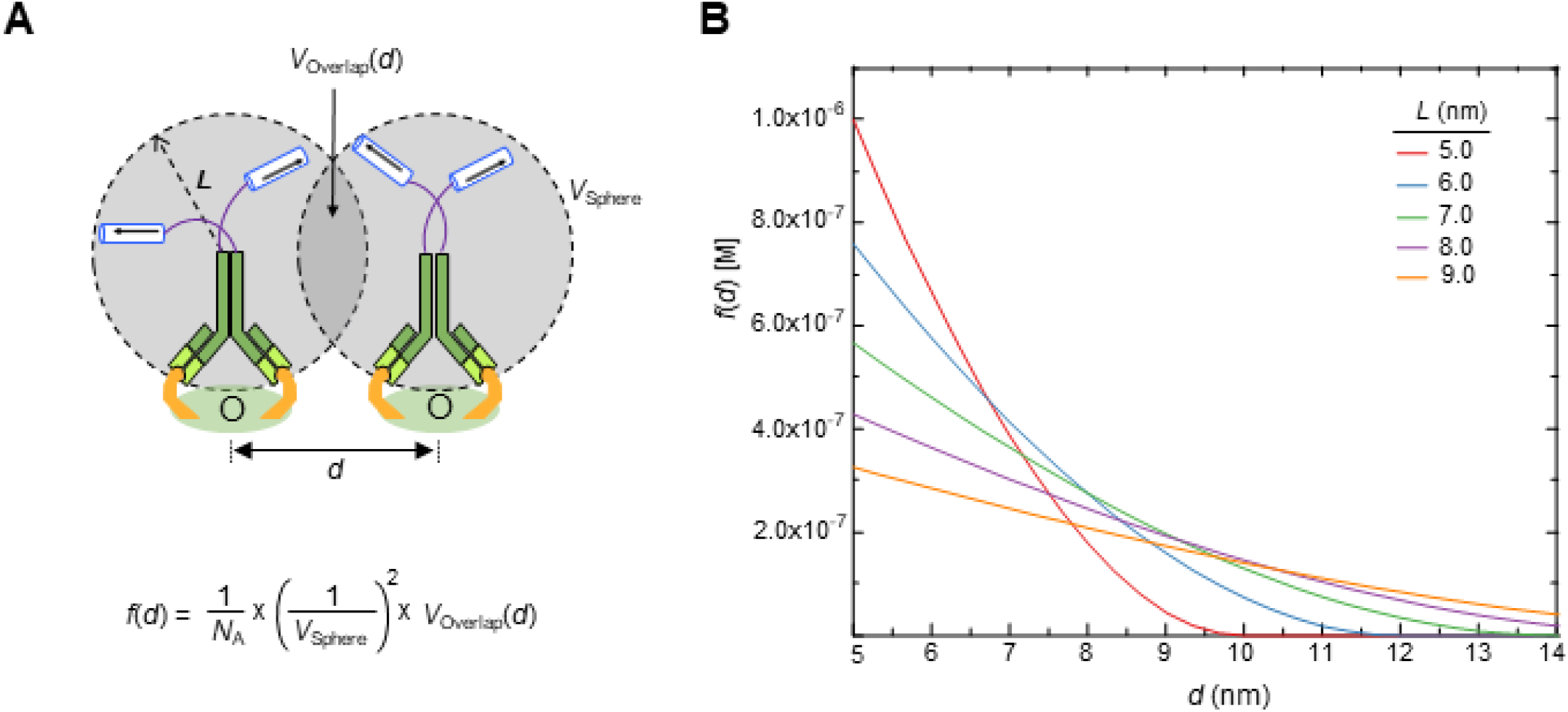
Calculation of *f*(*d*) using uniform local density approximation. The forward catenation rate at which two catenators dimerize is proportional to the volumetric overlap (*V*(*d*)) between the effective concentration of the catenator, which is assumed to be uniformly distributed over a sphere defined by the reach length (*L*). *V*(*d*) depends on the distance (*d*) between the two adjacent ^cat^Ab-2Ag complexes as well as the reach length. Shown on the left is a plot of *f*(*d*) as a function of *d* calculated for the indicated reach length (*L*).

**Figure 4—figure supplement 1.**
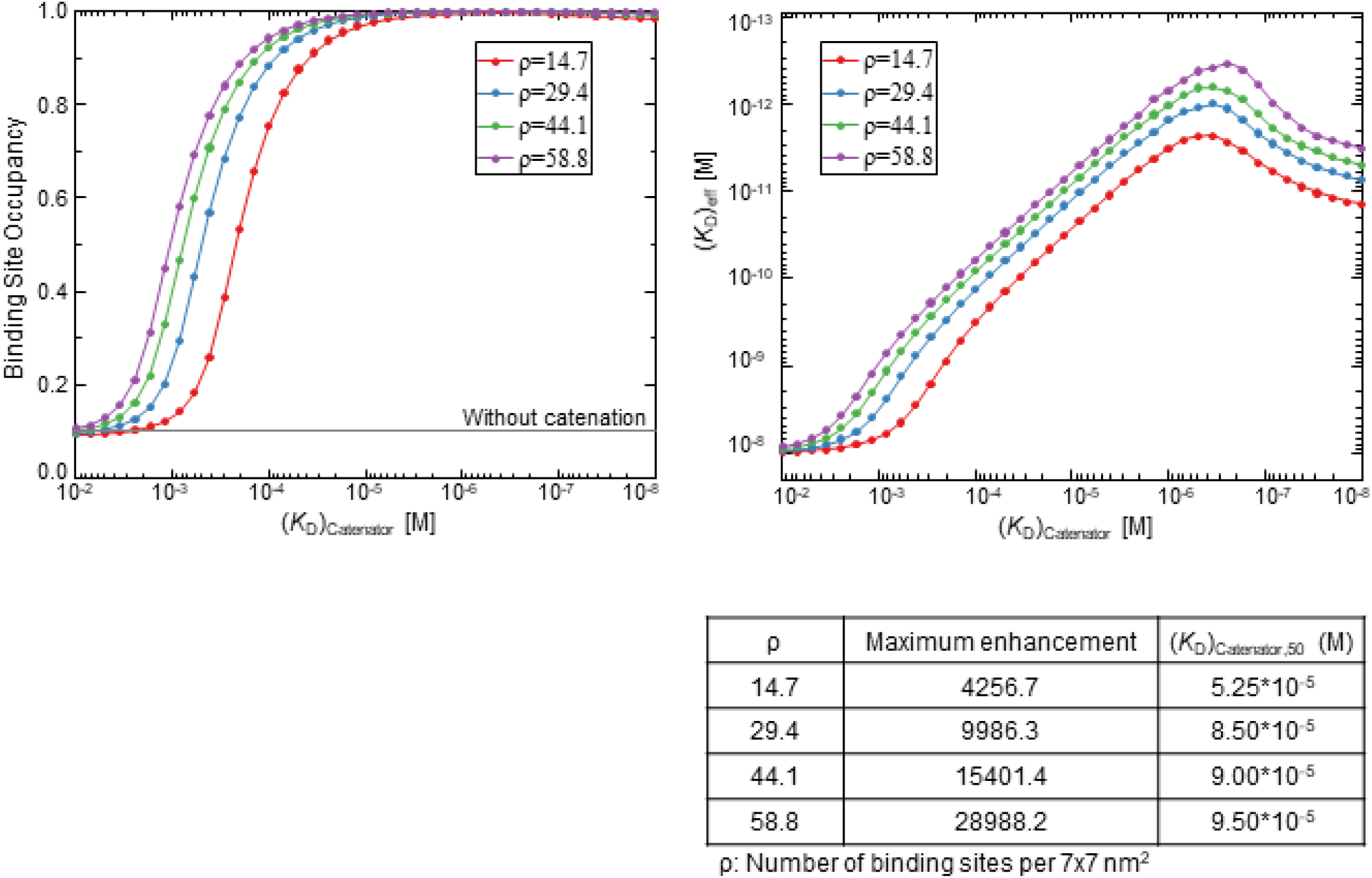
Simulations for randomly distributed, high-density binding sites. 1024 trials were sampled for each (*K*_D_)_catenator_ value at the indicated density and the results are plotted. The variables were *K*_D_= 10^-8^ M, [^cat^Ab]= 10^-9^ M, reach length= 7 nm, spacing between the binding sites= 12 nm, and the surface area= 5,760 nm^2^. The maximum fold enhancement of the effective binding avidity and (*K*_D_)_catenator,50_ are tabulated at the bottom.

**Figure 4—figure supplement 2.**
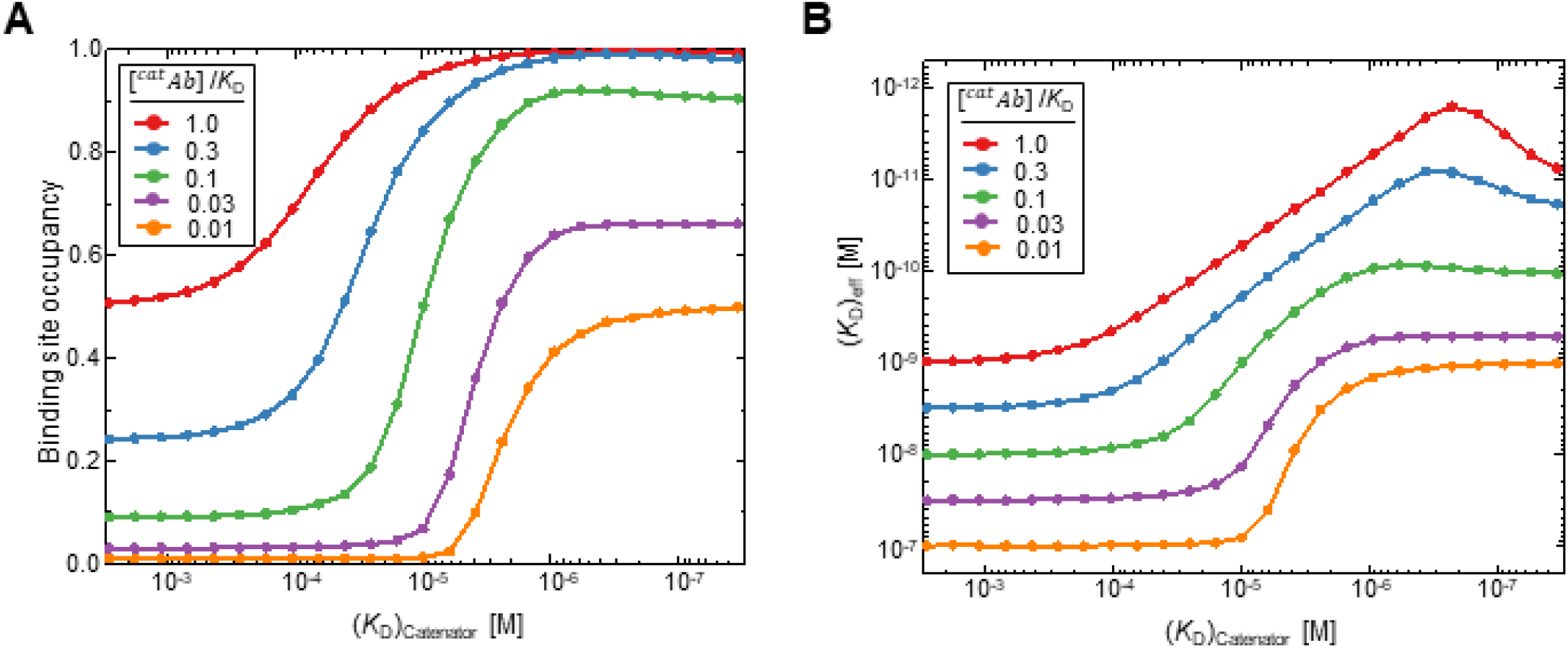
Influence of the likelihood of intrinsic antigen binding ([^cat^Ab]/*K*_D_) on binding site occupancy and (*K*_D_)_eff_. **(A)** The binding occupancy and (**B**) The effective dissociation constant (*K*_D_)_eff_ in response to [^cat^Ab]/*K*_D_ for [^cat^Ab]/*K*_D_ = 1.0, 0.3, 0.1, 0.03, 0.01. The simulations were carried out with a square array of the binding sites as in Figure 3. The set values for the variable parameters were [^cat^Ab]= 10^-9^ M, reach length= 7 nm, spacing between the binding sites= 12 nm and the number of total binding sites= 98. The mean binding occupancy of 1024 MCMC simulations was plotted. (*K*_D_)_catenator_ was varied from 3 mM to 30 nM. The antibody binding avidity is substantially enhanced across a broad range of [^cat^Ab]/*K*_D_.

